# High extracellular osmolarity promotes yeast thermotolerance through osmotic modulation and glycerol-dependent adaptation

**DOI:** 10.64898/2026.04.23.720099

**Authors:** Saki Oyama, Kosuke Shiraishi, Satoshi Okada, Fumiyoshi Abe, Anna Ahara, Reina Ito, Kokona Kosuga, Emiko Kusumoto, Tetsuo Tokiwano, Hiroya Yurimoto, Takashi Ito, Nobushige Nakazawa, Yuki Yoshikawa

**Affiliations:** Department of Biotechnology, Faculty of Bioresource Science, Akita Prefectural University, Akita 010-0195, Japan; Division of Applied Life Sciences, Graduate School of Agriculture, Kyoto University, Kyoto 606-8502, Japan; Department of Biological Science, Faculty of Science and Engineering, Yasuda Women’s University, Hiroshima 731-0153, Japan; Department of Biochemistry, Kyushu University Graduate School of Medical Sciences, Fukuoka 812-8582, Japan; Department of Chemistry and Biological Science, College of Science and Engineering, Aoyama Gakuin University, Kanagawa 252-5258, Japan

**Keywords:** Extracellular osmolarity, Thermotolerance, Glycerol metabolism, *Saccharomyces cerevisiae*, Hog1, Stress response

## Abstract

High-temperature stress is a major constraint on yeast growth and fermentation, and has traditionally been interpreted primarily in terms of intracellular molecular damage such as protein denaturation and aggregation. Despite this extensive focus on intracellular mechanisms, how physical factors within the extracellular environment influence yeast thermotolerance remains poorly understood. Here we demonstrate that increases in extracellular osmolarity markedly attenuate growth inhibition under high-temperature conditions in yeast. This protective effect was consistently observed in multiple laboratory and industrial strains of *Saccharomyces cerevisiae*, as well as in ascomycetous and basidiomycetous yeasts, indicating that osmotic pressure-dependent thermotolerance is a broadly conserved phenomenon. We also found that extracellular osmolarity dynamically increases during growth and then decreases in a diauxic shift-like pattern after growth arrest. At high temperature, the secretion of glucose-derived metabolites decreased, but that of other solutes increased, suggesting that heat stress alters the composition of extracellular solutes contributing to osmolarity. In addition, intracellular glycerol levels increased at high temperature, and this increase was further enhanced under high-osmolarity conditions. Notably, expression of a constitutively active Hog1 mutant exhibited raised intracellular glycerol levels, enhanced nuclear localization of Hog1, and improved growth under high-temperature conditions. Collectively, these findings support a model in which extracellular osmolarity is modulated to avoid excessive intracellular osmolarity under high-temperature conditions, while the intracellular accumulation of glycerol contributes to yeast adaptation at high-temperature. Our results highlight extracellular–intracellular osmotic coordination as an additional physiological layer of high-temperature stress adaptation in yeast.

## 1 Introduction

High-temperature stress is a major environmental challenge that limits cellular growth, viability, and productivity in microorganisms including yeast (Verghese et al., 2012; Zhou et al. 2021). In *Saccharomyces cerevisiae*, the effects of high temperature and thermotolerance mechanisms have been extensively investigated from both fundamental and applied perspectives (Techaparin et al. 2017; Yamamoto et al. 2008). A prevailing concept is that the deleterious effects of high-temperature stress are primarily due to molecular damage—particularly protein denaturation, misfolding, and aggregation, all of which disrupt essential cellular processes (Hänninen et al. 1999; Trotter et al. 2001; Yost et al. 1991; Shahsavarani et al. 2012; Ingolia & Craig 1982). Consistent with this view, numerous studies have focused on molecular chaperones, proteostasis networks, and quality control systems that mitigate thermal damage and restore intracellular homeostasis. The molecular damage-centric framework is further supported by genetic, evolutionary, and breeding studies of thermotolerant yeast strains (Huang et al. 2018; Shahsavarani et al. 2012). In addition to cytosolic proteins, high-temperature stress can affect membrane-associated proteins, potentially compromising the structural integrity of the plasma membrane and increasing its permeability (Godinho et al. 2020; Guyot et al. 2015), which in turn is strongly associated with cellular damage. Collectively, these findings have established protein denaturation and its downstream consequences as a primary source of cellular damage under high-temperature conditions.

Despite this strong emphasis on macromolecular damage, several observations suggest that non-macromolecular factors also contribute to thermotolerance in microorganisms. For example, Papouskova and Sychrova (2007) reported that osmotic supplementation with salt or sugar enhances thermotolerance in yeast cells. In addition, growth of yeast at high temperature has been shown to be improved in high-glucose medium (Costa et al. 2014). Furthermore, loss of the glycerol channel Fps1 results in increased heat sensitivity, which is alleviated by sorbitol addition, indicating that osmotic conditions critically influence yeast survival at high temperature (Beese et al. 2009). While these studies clearly demonstrate a phenotypic link between osmotic conditions and thermotolerance, the biophysical determinants (e.g., turgor pressure, membrane tension, or cell volume regulation) governing this interaction remain to be addressed.

More recently, Dunayevich *et al*. (2018) demonstrated that increased temperature activates the high osmolarity glycerol (HOG) signaling pathway via Fps1-mediated rapid efflux of glycerol, and proposed a model in which the reduction in turgor pressure caused by this abrupt glycerol efflux during temperature change triggers HOG signaling. Another study has shown that intracellular glycerol accumulation during growth under high temperature occurs independently of Hog1 (Siderius et al. 2000). This suggests that at least two pathways—differing in timing and mechanism—are involved in temperature-induced glycerol dynamics. In particular, the physiological significance of the long-term intracellular accumulation of glycerol during high-temperature growth, and its relationship with dynamic changes in the extracellular environment, remains poorly understood.

As well as signaling responses, high temperatures affect yeast cellular energy metabolism and cytoplasmic physical properties. At high temperature, proton extrusion mediated by the plasma membrane H⁺-ATPase increases, contributing to the maintenance of intracellular pH homeostasis (Coote et al. 1991). At the same time, prolonged high-temperature cultivation may lead to ATP depletion, triggering the synthesis and accumulation of storage carbohydrates such as trehalose and glycogen (Persson et al. 2020). These metabolic changes are closely linked to the physical properties of the cytoplasm. In general, increasing temperature decreases the viscosity of aqueous solutions and accelerates the diffusion of particles (Longsworth, 1954); however, yeast cells can counteract this effect through a process termed “viscoadaptation”, in which the carbohydrates that are accumulated during high-temperature cultivation increase intracellular viscosity (Persson et al. 2020). To date, viscoadaptation has been attributed mainly to the accumulation of trehalose and glycogen, and the potential contribution of other osmolytes has received little attention. In particular, it remains unclear whether glycerol—a major compatible solute in yeast that is known to accumulate in response to osmotic stress—contributes to the regulation of cytoplasmic physical properties during high-temperature growth.

From a physical perspective, extracellular osmolarity determines the osmotic gradient across the cell boundary and thereby influences turgor pressure, a key mechanical parameter affecting cell expansion, membrane tension, and structural integrity (Atilgan et al. 2015; Minc et al. 2014; Gervais et al. 1996; Mishra et al. 2022). Yeast cells possess well-characterized mechanisms to adapt to increases in extracellular osmolarity, prominently involving the synthesis and retention of glycerol as a compatible solute (Blomberg 2022; Schaber et al. 2010; Mochizuki et al. 2023). Glycerol contributes to osmotic balance and stabilizes proteins because it is preferentially excluded from a protein’s surface, thereby enhancing hydration and rendering the denatured state thermodynamically unfavorable, which in turn provides broad protection against physicochemical stress (Meng et al. 2004; Panadero et al. 2006; Gekko et al. 1981; Pocivavsek et al. 2011). A recent review has emphasized the multilayered regulation of glycerol metabolism and the existence of regulatory routes beyond the canonical HOG signaling pathway (Blomberg 2022). While previous studies have provided extensive knowledge on osmotic stress responses and HOG-mediated regulation, much of our understanding is built upon studies of acute stress responses, particularly in the context of rapid osmotic shifts. Consequently, the contribution of the HOG signaling pathway and related regulatory networks to glycerol accumulation and sustained growth during prolonged cultivation under high-temperature conditions remains largely unresolved.

To address the above gap in our knowledge, here we have explored the interplay among extracellular osmolarity, glycerol metabolism, and growth phenotypes under prolonged high-temperature conditions in yeast. We have investigated growth-dependent changes in extracellular osmolarity under optimal and high-temperature conditions and examined their impact on yeast thermotolerance. Through the integration of extracellular environment measurements with analyses of glycerol dynamics, we are able to propose a mechanism for regulating intracellular and extracellular osmotic pressure via solute efflux under high-temperature conditions. Our findings suggest that high extracellular osmolarity helps to buffer the excessive osmotic gradient across the cell boundary that occurs during growth at high temperature. Furthermore, we have identified glycerol-dependent adaptation as a key mechanism contributing to growth under high-temperature stress.

## 2 Materials and Methods

### 2.1 Yeast strains and media

The yeast strains and primers used in this study are listed in Tables 1 and 2, respectively. *S. cerevisiae* BY4741 (*MATa his3*Δ*1 leu2*Δ0 *met15*Δ0 *ura3*Δ0) was used as the wild-type (WT) strain and as the host strain for strain construction. In the BY4741 background, we generated *fps1*Δ0 and *hog1*Δ0 deletion mutants, as well as a constitutively active Hog1 variant (Hog1^F318L^) (Bell et al. 2001), by genome editing using a previously described method (Okada et al. 2021) with minor modifications. The genome editing primers and plasmids are listed in Tables 2 and 3, respectively.

**TABLE 1.** *S. cerevisiae* strains used in this study.

| Strain name | Strain number | MAT | Relevant genotype | Strain background | sgRNAs/crRNAs | Reference/Resource |
| --- | --- | --- | --- | --- | --- | --- |
| BY4741 | YIT5001 | a | *his3Δ1 leu2Δ0 met15Δ0 ura3Δ0* | BY4741 | - | Brachmann et al. 1998 |
| YIT16090 | YIT16090 | a | *hog1Δ0* | BY4741 | HOG1Sp3 | This study |
| YIT16383 | YIT16383 | a | *hog1(F318L)* | BY4741 | HOG1Sp3 | This study |
| YIT16631 | YIT16631 | a | *fps1Δ0* | BY4741 | FPS1Sp3 | This study |
| WT-Hog1-GFP | AYS001 | a | *HOG1-EGFP* | BY4741 | - | This study |
| F318L-Hog1-GFP | AYS002 | a | *hog1(F318L)-EGFP* | YIT16383 | - | This study |
| S288C | BY611 | α | *SUC2 mal mel gal2 CUP1 flo1 flo8-1 SSD1-v1* | - | - | Goffeau et al. 1996/NBRP |
| W303 | BY4502 | a/α | *ade2-1/ade2-1 his3-11,15/his3-11,15 leu2-3,112/leu2-3,112 trp1-1/trp1-1 ura3-1/ura3-1 can1-100/can1-100* | - | - | Markus Ralser, 2012/NBRP |
| X2180 | BY4947 | a/α | *SUC2 mal mel gal2 CUP1 flo1 flo8-1 SSD1-v1/SUC2 mal mel gal2 CUP1 flo1 flo8-1 SSD1-v1* | - | - | Mortimer & Johnston 1986/NBRP |
| Σ1278b | BY29040 | α | wild type *MPR1 MPR2 AZCr* | - | - | Dowell et al. 2010/NBRP |
| SK1 | 204722 | a/α | *can1(r) gal2 cup(s)* | - | - | Börner & Cha 2015/ATCC |
| K7 | RIB1003 | a/α | *msn2 msn4 AWA1* | - | - | Watanabe et al. 2011/RIB |
| K701 | RIB1004 | a/α | *msn2 msn4 awa1* | RIB1003 | - | Miyashita et al. 2004/RIB |

**TABLE 2.**
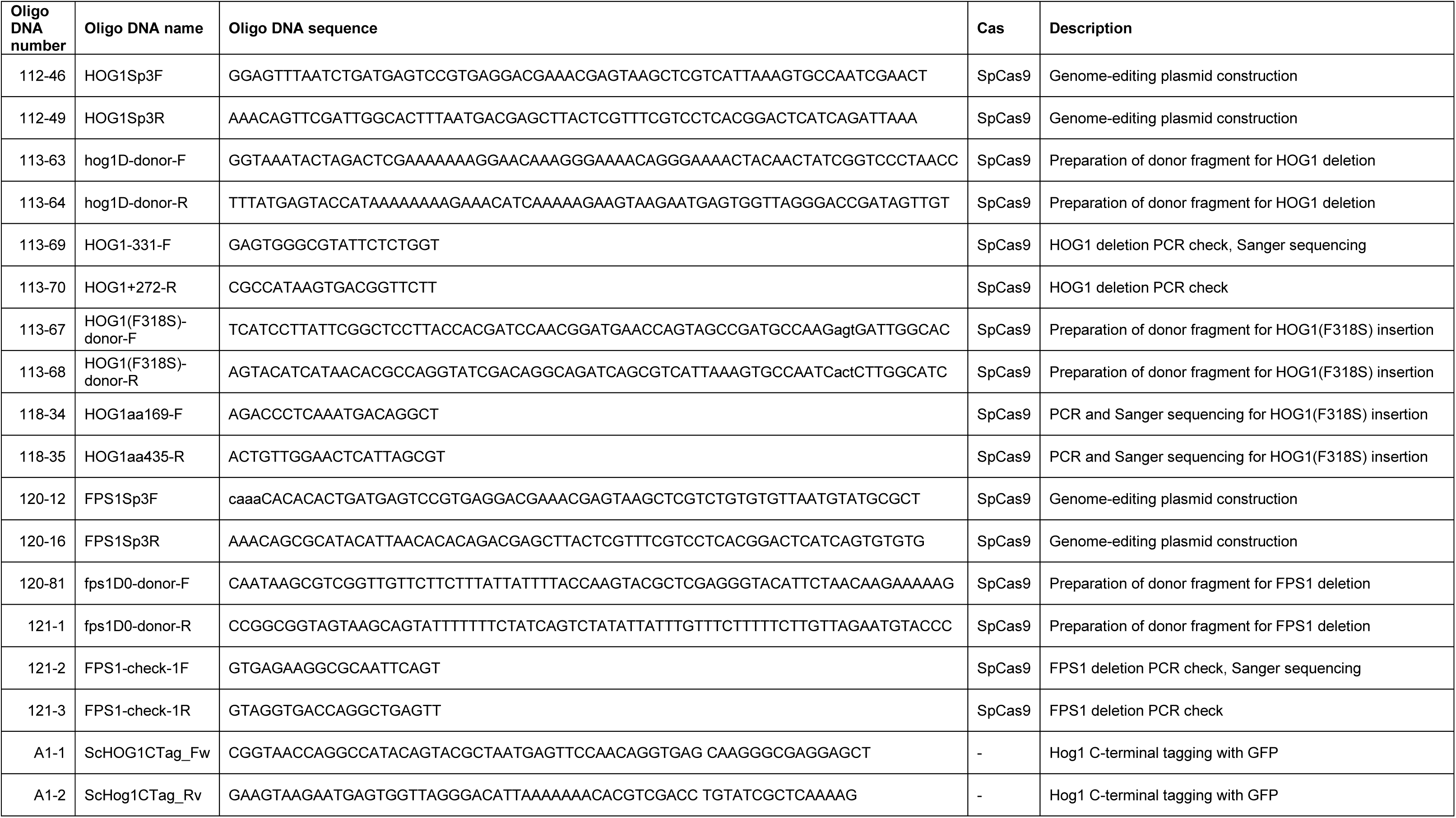
Oligo DNA sequences used in this study.

**TABLE 3.**
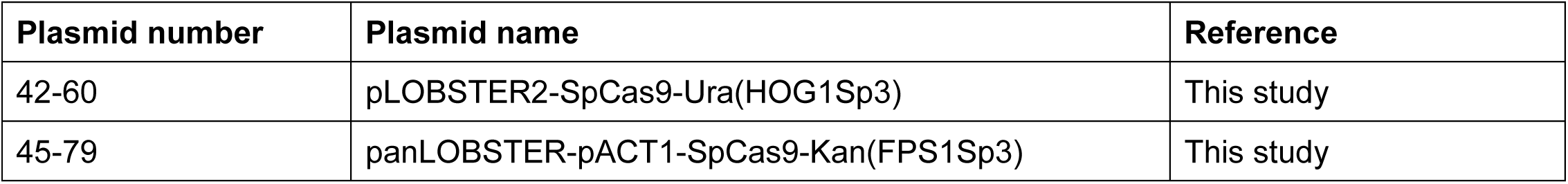
Genome-editing plasmids used in this study.

Each genome-editing plasmid encoded SpCas9 fused to an SV40 nuclear localization signal and a guide RNA (gRNA); expression of these components was driven by either the *GAL1* or *ACT1* promoter. gRNAs were designed using CRISPOR (Haeussler et al. 2016). PCR-generated donor fragments containing homology arms flanking the cleavage site were used to introduce deletions or an insertion.

Host cells were co-transformed with the genome-editing plasmid and the donor fragment, spread on agar plates containing SC−Ura medium supplemented with 2% galactose or YP supplemented with 2% galactose and 200 μg/mL of G418 (Nacalai Tesque), and incubated at 30 °C for 3–5 days. The resulting colonies were picked and re-streaked on fresh plates. Successful editing was screened by PCR across the target locus and confirmed by Sanger sequencing of the PCR product. The cells were then grown in liquid YPD to remove the genome-editing plasmids. Single colonies were re-streaked on YPD and on SC (Dex)−Ura or YPD + G418 plates, and clones confirmed to have lost the editing plasmid were selected and stored for subsequent experiments.

To construct strains expressing GFP-tagged Hog1 in WT and *hog1*-F318L cells, DNA fragments were amplified by PCR using plasmid pMO152 as a template and primers ScHog1CTag_Fw and ScHog1CTag_Rv (Shiraishi et al. 2018), and then integrated into the C-terminal region of the *HOG1* gene in either BY4741 or *hog1*-F318L cells expressing Hog1-GFP or F318L-Hog1-GFP, respectively.

The other laboratory strains, S288C, X2180 (X2180-1A×X2180-1B), W303, Σ1278b, and SK1, were obtained from the National BioResource Project (NBRP) or the American Type Culture Collection. Industrial sake yeast strains Kyokai no. 7 (K7) and no. 701 (K701) were provided by the National Research Institute of Brewing, Japan (Table 1). The non-*Saccharomyces* ascomycetous and basidiomycetous yeast strains used in this study (Table 4) were obtained from the NBRP and NITE Biological Resource Center.

**TABLE 4.**
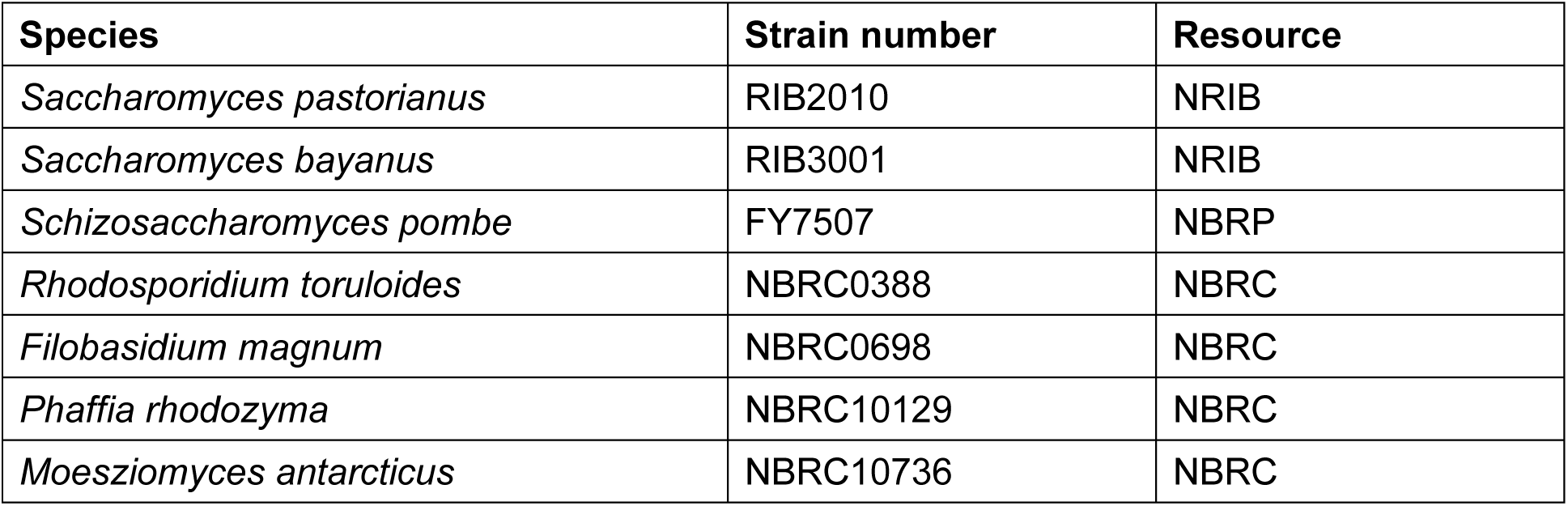
Non-*S. cerevisiae* strains used in this study.

Yeast cells were cultured at 30 °C in yeast extract–peptone–dextrose (YPD) medium containing 1% (wt/vol) yeast extract, 2% (wt/vol) peptone, and 2% (wt/vol) glucose with 2% (wt/vol) agar if necessary. For high-osmolarity conditions, sorbitol or KCl was added to the medium at the indicated concentrations.

### 2.2 Assessment of cell growth and correlation between OD_600_ and biomass at each temperature

Cell growth in liquid cultures was monitored by measuring optical density at 600 nm (OD_600_). For selected experiments, dry cell weight (DCW) was determined to assess the relationship between OD_600_ and biomass at different temperatures. In brief, defined volumes of culture were harvested, washed, and dried at 100 °C until a constant weight was achieved.

For qualitative assessment of growth phenotypes across yeast species and strains, spot assays were performed. Serial dilutions of cell suspensions were spotted onto solid YPD agar plates supplemented as indicated and incubated at the designated temperatures.

### 2.3 Phylogenetic analysis

Phylogenetic analysis was performed based on the D1/D2 domain of the large subunit rRNA gene. Representative LSU D1/D2 sequences for each species were retrieved from the GenBank database. When strain-specific sequences were unavailable, sequences from type strains or well-characterized reference strains of the same species were used.

The following accession numbers were used for phylogenetic tree construction: *S. cerevisiae* (KU837254.1), *S. pastorianus* (AY048172.1), *S. bayanus* (AF113892.1), *Schizosaccharomyces pombe* (KC111877.1), *Rhodosporidium toruloides* (IF00038801), *Filobasidium magnum* (IF00069801), *Phaffia rhodozyma* (IF01012901), and *Moesziomyces antarcticus* (AJ235302.1).

Sequences were aligned using MAFFT (Katoh, 2013) with default parameters. Poorly aligned terminal regions containing gaps or missing data were trimmed, retaining only alignment positions shared across all taxa. A phylogenetic tree was constructed using IQ-TREE 3.0.1 (Minh et al. 2020; Wong et al. 2025) with ModelFinder (Kalyaanamoorthy et al. 2017). Branch support was assessed by ultrafast bootstrap analysis with 1000 replicates. The resulting tree was visualized using iTOL (Letunic & Bork 2021).

### 2.4 Normalization of biomass using OD_600_ and DCW

Optical density at 600 nm was used as an initial proxy for cell density. Because biomass per unit OD₆₀₀ was found to vary by cultivation temperature, OD_600_ values were corrected using temperature-specific conversion factors derived from DCW measurements (see Section 2.2). For each cultivation temperature (30 °C, 37 °C, 39 °C, and 40 °C), the relationship between OD_600_ and DCW was determined from more than 10 independent biological samples (Figure S3). DCW per unit OD_600_ was calculated for each condition, and values were normalized to those obtained at 30 °C to derive correction factors.

These temperature-specific correction factors were applied throughout the study to normalize OD-based measurements, including extracellular osmolarity or glycerol, and intracellular glycerol quantification (Figures 2 and 3), thereby ensuring accurate comparison on a biomass-equivalent basis.

**FIGURE 1.**
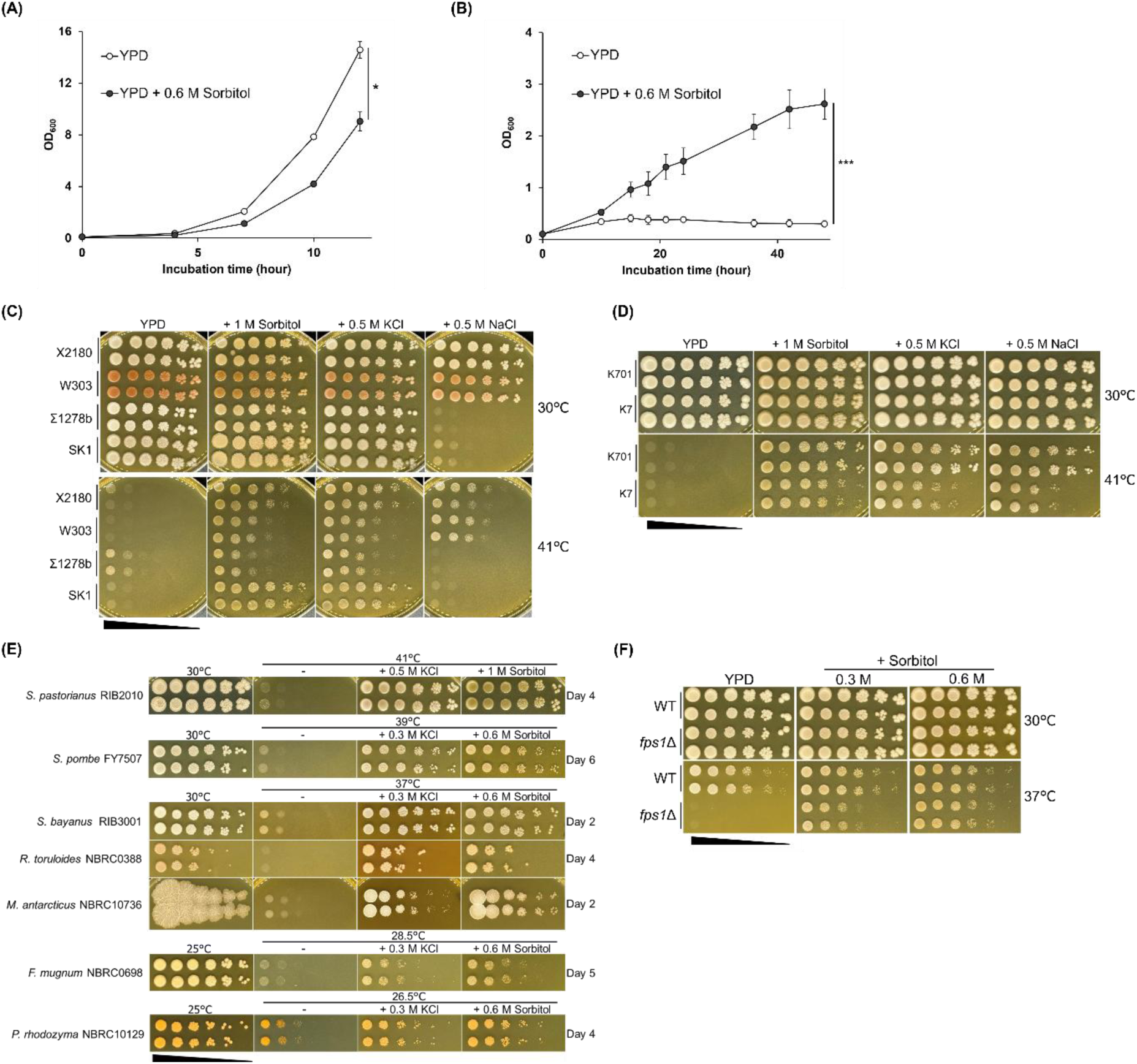
Increased extracellular osmolarity alleviates high temperature–induced growth inhibition in yeast. (A, B) Growth of strain S288C in YPD medium with or without 0.6 M sorbitol at 30 °C (A) and 42 °C (B). (C–E) Spot assay analysis of growth at high temperature with or without osmotic supplementation in laboratory and industrial yeast strains. (F) Growth of *fps1*Δ cells at high temperature in the presence or absence of sorbitol supplementation. Data are the mean ± SD of at least three independent biological replicates. Statistical significance was determined by repeated-measures ANOVA (sorbitol × time interaction); \**p* < 0.05, \*\*\**p* < 0.001.

**FIGURE 2.**
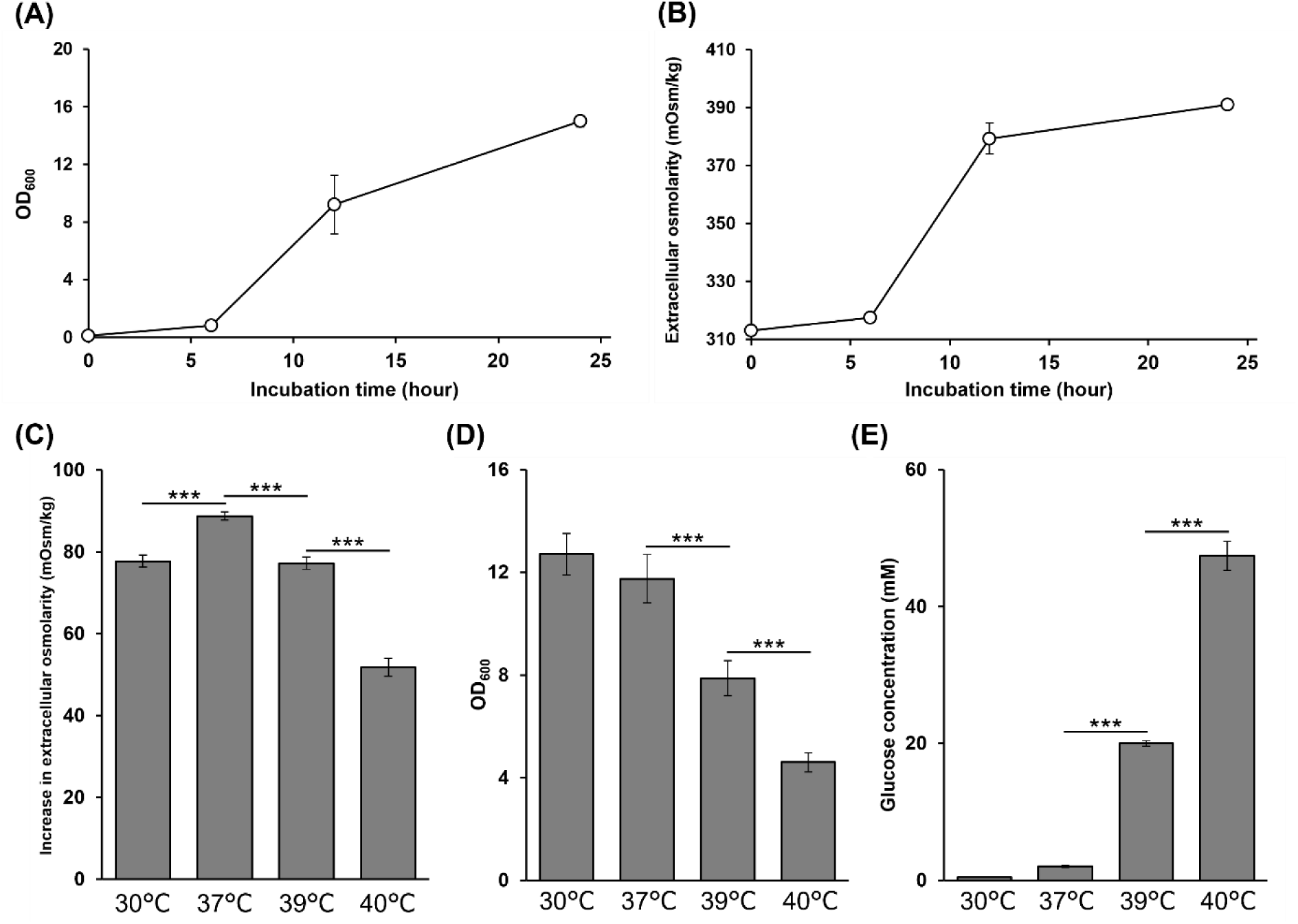
Temperature-dependent changes in extracellular osmolytes, cell growth, and residual glucose levels during yeast cultivation. (A, B) Growth of WT BY4741 in YPD medium at 30 °C (A) and extracellular osmolarity of the culture supernatant (B). (C) Increase in extracellular osmolarity in the culture supernatant after growth at the indicated temperatures. The increase in osmolarity was calculated by subtracting the osmolarity of the original YPD medium from that of the culture supernatant after cultivation. (D) Cell growth, measured as OD_600_, after cultivation at the indicated temperatures. Because the relationship between OD_600_ and biomass varied depending on cultivation temperature, OD_600_ values were corrected using the experimentally determined relationship between OD_600_ and dry cell weight (Figure S3). (E) Residual glucose concentrations in the same culture supernatants used for osmolarity measurements. Data are the mean ± SD of at least three independent biological replicates. Statistical significance was determined by one-way ANOVA with Tukey’s post-hoc test; \*\*\**p* < 0.001

**FIGURE 3.**
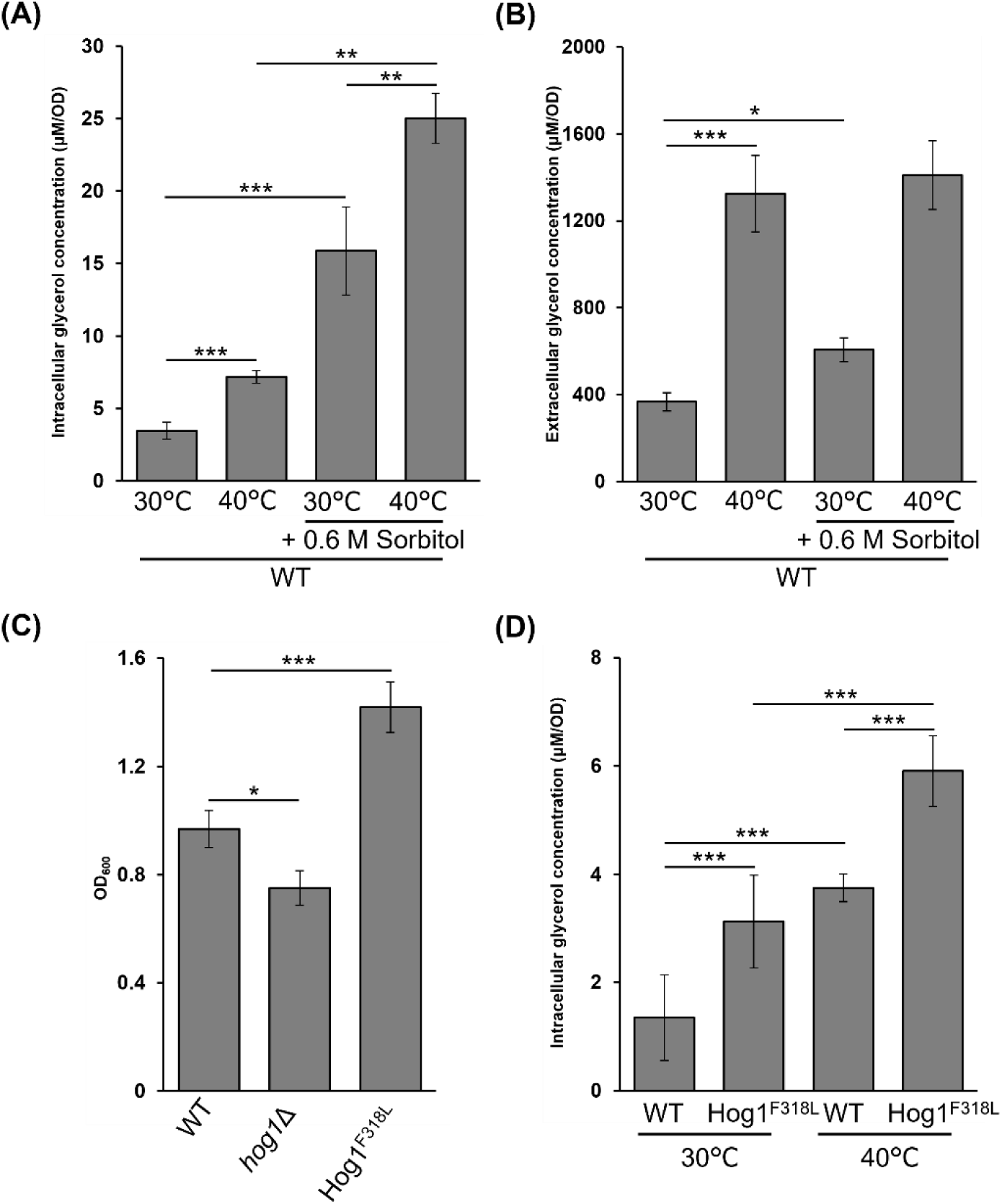
Temperature-dependent glycerol accumulation and its association with HOG pathway activity. For samples cultivated at high temperature, OD_600_ values were corrected using temperature-specific conversion factors derived from dry cell weight measurements (Figure S3), and glycerol levels were normalized on a biomass-equivalent basis. (A, B) Respective intracellular and extracellular glycerol levels in WT cells incubated under the indicated temperature and osmotic conditions for 16 h. (C) Growth of WT and Hog1^F318L^ mutant cells after cultivation at 41 °C for 24 h, assessed by OD_600_. (D) Intracellular glycerol levels in WT, *hog1*Δ, and Hog1^F318L^ mutant cells after incubation at the indicated temperature for 18 h. Data are the mean ± SEM of at least three independent experiments. Statistical significance was determined by two-way ANOVA (main effects of temperature and sorbitol, and their interaction) for (A), (B), and (D), and by one-way ANOVA with Tukey’s post-hoc test for (C); \**p* < 0.05, \*\**p* < 0.01, \*\*\**p* < 0.001.

### 2.5 Measurement of extracellular osmolarity

At designated time points, culture samples were collected and immediately centrifuged to remove cells. Supernatants were further clarified by filtration through a 0.45-μm membrane filter (DISMIC 03CP045AN) to ensure complete removal of residual cells. Extracellular osmolarity was measured using a micro-osmometer (Micro Osmometer OM-819, Bio Medical Science). To enable comparison across conditions with differing growth rates, osmolarity values were normalized to cell density at the time of sampling. Where OD_600_ did not accurately reflect biomass under high-temperature conditions, correction factors derived from OD_600_–DWC calibration experiments were applied.

### 2.6 Quantification of intracellular glycerol

For intracellular glycerol measurements, cell pellets were washed once with ultrapure water and resuspended in ultrapure water. The OD_600_ of the suspension was recorded for subsequent normalization. Aliquots (1 mL) were transferred to sealed microcentrifuge tubes and subjected to heat extraction at 100 °C for 30 min. After cooling on ice, samples were centrifuged at low temperature, and the supernatants were collected.

To remove proteins, three volumes of pre-chilled acetonitrile (LC–MS grade) were added to each sample, followed by incubation at −20 °C. After centrifugation, the supernatants were filtered through a hydrophilic PTFE membrane (0.22 μm; Shimadzu GLC) and subjected to liquid chromatography–tandem mass spectrometry (LC–MS/MS) analysis.

### 2.7 Quantification of extracellular glycerol

Extracellular glycerol was quantified from filtered culture supernatants prepared as described above. Samples were processed in parallel with intracellular extracts to ensure consistency. LC–MS/MS analysis was performed under identical chromatographic and detection conditions.

### 2.8 Liquid chromatography–tandem mass spectrometry

Liquid chromatography was performed on a Dionex UltiMate 3000 system (Thermo Fisher Scientific, MA, USA) equipped with a TSKgel OH-120 column (4.6 mm × 150 mm, 5 µm; Tosoh, Tokyo, Japan). The column oven temperature was maintained at 35 °C. The mobile phase consisted of 10% acetonitrile in water containing 10 mM ammonium acetate, delivered at a flow rate of 1.5 mL/min. The LC eluent was split using a T-splitter before entering the mass spectrometer; the flow rate introduced into the electrospray ionization (ESI) source was reduced to 0.2 mL/min.

Mass spectrometry was conducted using TSQ Quantum Ultra triple quadrupole mass spectrometer (Thermo Fisher Scientific, MA, USA) equipped with an ESI source. The spectrometer was operated in negative ionization mode using selected ion monitoring. Target analytes were detected as acetate adduct ions ([M + CH_3_COO]^−^) at *m/z* 151.0 for glycerol and *m/z* 156.0 for glycerol-d_5_. The ESI source parameters were set as follows: ionization voltage, 3000 V; capillary temperature, 300 °C. Data acquisition and processing were performed using Xcalibur software (version 4.0, Thermo Scientific).

Quantification was performed by using glycerol-d_8_ (Sigma-Aldrich 447498) as an internal standard. Due to the rapid H/D exchange of the three hydroxyl deuterons with the protic solvent, the internal standard was detected and monitored as the [1,1,2,3,3-d_5_] glycerol isotopologue (*m/z* 156.0, as an acetate adduct). Notably, the signal intensities for ions corresponding to isotopologues were less than 1% of that of the species, indicating that the exchange was essentially complete under the analytical conditions. Calibration curves were constructed by plotting the peak area ratio of glycerol (*m/z* 151.0) to glycerol-d_5_ (*m/z* 156.0) against the nominal concentration of glycerol. Linear regression analysis was used to fit the data, and the curves showed excellent linearity with a correlation coefficient (r^2^) greater than 0.99 over the concentration range of 6.85 to 274 µM.

### 2.9 Hog1 subcellular localization under acute stress conditions

Subcellular localization of Hog1 was analyzed using the *hog1*Δ strain expressing EGFP-tagged WT Hog1 (WT-Hog1-EGFP) or the constitutively active Hog1^F318L^ variant (Hog1-F318L-EGFP). Cells were pre-cultivated in YPD medium for 24 hours and transferred to synthetic defined (SD) medium (0.67% Yeast Nitrogen Base without amino acids, 2% glucose). Cultures were then subjected to acute stress treatments. After hyperosmotic stress (0.6 M sorbitol) or high-temperature stress (40 °C), cells were collected at the indicated time points.

Fluorescence microscopy was performed using an IX83 inverted fluorescence microscope (Evident, Tokyo, Japan) equipped with an IX3-SSU motorized stage (Evident), a drift compensation module (IX3-ZDC, Evident), and a filter wheel unit (96A357 and MAC6000, Ludl Electronic Products Ltd., Hawthorne, NY, USA) suitable for detecting fluorescent proteins. Images were captured with an ORCA-Fusion Digital CMOS camera (USB3.0 set, Hamamatsu Photonics, Hamamatsu, Japan) and analyzed by cellSens software (Evident) and Fiji.

### 2.10 Nuclear staining

After cultivation for the indicated time, cells were harvested, washed once, and fixed with 1 mL of 70% ethanol for 30 min at room temperature. Fixed cells were washed twice, resuspended in 150 µL of sterilized water, and stained with 150 µL of 0.125 µg/mL of 4′,6′-diamidino-2-phenylindole (DAPI) solution. After 10 min of incubation, the samples were visualized by microscopy.

### 2.11 Statistical analysis

All statistical analyses were performed using EZR (Jichi Medical University, Tochigi, Japan)(Kanda 2013), a modified version of R Commander (The R Foundation for Statistical Computing, Vienna, Austria) with additional functions frequently used in biostatistics. All experiments were performed with at least three independent biological replicates. Differences among multiple conditions were evaluated by using analysis of variance (ANOVA), followed by appropriate post hoc tests when applicable. Statistical significance was defined as \**p* < 0.05, \*\**p* < 0.01, and \*\*\**p* < 0.001.

## 3 Results

### 3.1 High extracellular osmolarity attenuates heat-induced growth inhibition in yeast

To examine the effect of extracellular osmolarity on yeast growth under high-temperature conditions, growth was first assessed in liquid culture using the laboratory type strain *S. cerevisiae* S288C (Goffeau et al. 1996) (Figure 1A, B). At the optimal temperature (30 °C), high extracellular osmolarity impaired growth, consistent with growth inhibition caused by osmotic stress, as previously described (Altenburg et al. 2019; Watson 1970). In contrast, at 42 °C, which caused severe inhibition of growth under standard conditions, supplementation with sorbitol partially restored yeast growth in a concentration-dependent manner.

To determine whether this protective effect depends on genetic background, we performed spot assays using the following *S. cerevisiae* laboratory and industrial strains with distinct genetic features: X2180, a derivative of the strain S288C background (Mortimer & Johnston 1986)); W303, derived from strain S288C but exhibiting numerous differences, including amino acid synthesis (Markus Ralser, 2012); Ʃ1278b, a filamentous-growth–competent strain genetically divergent from strain S288C (Dowell et al. 2010); SK1, a diploid strain widely used for studies of meiosis and sporulation (Börner & Cha 2015); and the industrial sake strains K7, which lacks Msn2 and/or Msn4 stress-response regulators (Watanabe et al. 2011), and the nonfoaming mutant K701 (Miyashita et al. 2004)(Figure 1C, D). Across these diverse strains, high extracellular osmolarity reproducibly alleviated growth defects under high-temperature conditions, indicating that the protective effect is not limited to a specific laboratory background. Similar growth restoration was observed with both non-ionic (sorbitol) and ionic (KCl and NaCl) osmolytes, indicating that the effect is primarily attributable to osmolarity rather than solute-specific chemical properties. Of note, Σ1278b and SK-1 exhibited marked NaCl sensitivity regardless of temperature, consistent with a previous report (David Engelberg, 2025), suggesting that strain-specific ion sensitivity may override the growth-restoring effect of high osmolarity observed at high temperature.

To further evaluate the generality of this effect across species, spot assays were extended to phylogenetically diverse yeasts including the ascomycetous species *S. bayanus*, *S. pastorianus*, and *Schizosaccharomyces pombe*, and the basidiomycetous species *Rhodosporidium toruloides*, *Filobasidium magnum*, *Phaffia rhodozyma*, and *Moesziomyces antarcticus*, spanning distinct ecological niches and evolutionary lineages (Figure 1E; Figure S1). Although the magnitude of the response varied among species, osmotic supplementation generally improved growth under high-temperature conditions across these evolutionarily and physiologically diverse yeasts. The exceptions were *F. magnum* and *P. rhodozyma*, which preferentially grow at low temperatures; these species failed to grow at temperatures above 30 °C even under high extracellular osmolarity conditions, and the osmotic growth-promoting effect was particularly limited in *P. rhodozyma*.

To evaluate the impact of aquaglyceroporin function deficiency on growth at high temperature, we used spot analysis to compare *FPS1* knockout and WT strains of BY4741, derived from strain S288C (Brachmann et al. 1998). The *fps1*Δ mutant showed strong growth inhibition at 37 °C, which was partially alleviated by supplementation with 0.3 M sorbitol and almost completely alleviated by 0.6 M sorbitol (Figure 1F). These results indicated that the osmotic pressure-mediated attenuation of growth inhibition at high temperatures is concentration-dependent.

### 3.2 A non-fermentation-derived source potentially increases extracellular osmolarity during high-temperature growth

Next, we characterized changes in extracellular osmolarity during yeast growth by monitoring the osmolarity of culture supernatants for WT BY4741 (Figure 2A, B). Extracellular osmolarity increased in parallel with cell growth, but then gradually decreased after reaching a peak. The decrease was cell-dependent and not observed in cell-free controls (Figure S2A, B). These results indicate that the total amount of substances excreted by yeast cells during growth exceeds the amount of nutrients taken up from the medium. The subsequent cell-dependent decrease in osmolarity is consistent with metabolic remodeling associated with the diauxic shift (Zampar et al. 2013; Brauer et al. 2005).

To assess the effect of temperature, extracellular osmolarity was measured after 15 hours of cultivation at the indicated temperatures (Figure 2C). The increase in osmolarity was enhanced at 37 °C, comparable at 30 and 39 °C, and reduced at 40 °C. Growth was significantly impaired at ≥39 °C (Figure 2D), and residual glucose levels increased under these conditions (Figure 2E), indicating reduced glucose consumption. Because OD600_nm_ did not equivalently reflect biomass at different temperatures, we subsequently corrected OD600_nm_ values based on the relationship between OD600_nm_ and DCW (Figure S3).

Despite the expected decrease in fermentation-derived products such as ethanol at high temperature, the magnitude of the increase in extracellular osmolarity remained comparable at 30 and 39 °C. This observation suggests that solutes other than those derived from glucose metabolism contribute to the increase in extracellular osmolarity at high temperature.

### 3.3 High temperature and osmotic stress independently and additively increase intracellular glycerol levels

Given the observed temperature-dependent changes in solute release during cultivation (Figure 2), we examined the levels of glycerol, a major compatible solute involved in osmotic regulation in yeast. Intracellular glycerol levels were quantified under different cultivation conditions and compared with those observed at the optimal growth temperature of 30 °C (Figure 3A,B).

Growth at high temperature resulted in a significant increase in intracellular glycerol levels relative to growth at the optimal temperature (Figure 3A). Similarly, at the optimal growth temperature, osmotic supplementation led to higher intracellular glycerol levels as compared with the no-supplementation control. When cells were subjected to both high temperature and osmotic supplementation, intracellular glycerol levels were higher than those observed under either condition alone. Notably, the combined treatment resulted in an approximately additive increase in glycerol accumulation as compared with either condition alone.

To determine whether the changes in intracellular glycerol were accompanied by alterations in extracellular glycerol, extracellular glycerol levels were also quantified under the two conditions (Figure 3B). Under high osmolarity, extracellular glycerol levels remained relatively low, consistent with intracellular glycerol accumulation. In contrast, high-temperature cultivation was associated with increased extracellular glycerol levels. These results indicate that high osmolarity conditions are accompanied by glycerol synthesis and accumulation, and that high temperatures promotes both glycerol synthesis and its efflux.

### 3.4 Modulation of the HOG signaling pathway alters glycerol accumulation and thermotolerance phenotypes

Lastly, we assessed the contribution of the HOG signaling pathway to glycerol accumulation and growth at high temperature by generating two yeast strains with altered *HOG1* activity (Figure 3C, D, Figure S4): a deletion mutant (*hog1*Δ), and a strain expressing a constitutively active Hog1 variant (Hog1^F318L^). The two strains were cultivated at various temperatures, and their growth phenotypes and intracellular glycerol levels were quantified.

Deletion of *HOG1* slightly reduced growth at high temperature as compared with the WT strain. Intracellular glycerol concentrations did not differ significantly between the *hog1*Δ mutant and WT at either the optimal temperature or high temperature, and glycerol concentrations in both strains were increased at high temperature relative to the optimal temperature (Figure S4). Thus, temperature-dependent glycerol accumulation is largely maintained in the absence of *HOG1*.

In contrast, the Hog1^F318L^ strain with constitutive activation of Hog1 exhibited higher intracellular glycerol concentrations relative to the WT strain at both the optimal temperature and high temperature (Figure 3C). Under high-temperature cultivation, the Hog1^F318L^ strain also showed improved growth relative to the WT strain (Figure 3D). These results suggest that enhanced HOG pathway activity is associated with elevated intracellular glycerol levels and improved growth at high temperature.

To investigate how constitutive activation of Hog1 may influence cellular stress responses, we examined the subcellular localization of WT Hog1 and the constitutively active Hog1^F318L^ variant by using the *hog1*Δ strain expressing WT-Hog1-EGFP or F318L-Hog1-EGFP. Under control conditions, both WT-Hog1-EGFP and F318L-Hog1-EGFP were predominantly distributed throughout the cytoplasm (Figure 4). Upon exposure to osmotic stress (0.6 M sorbitol), nuclear accumulation of Hog1 was induced in the WT-Hog1-EGFP-expressing strain, as indicated previously (Shiraishi et al. 2018). Notably, under the same conditions, a substantially higher proportion of cells in the F318L-Hog1-EGFP-expressing strain showed Hog1 localization to a specific subcellular compartment (Figure 4A). This compartment was confirmed to be the nucleus by co-staining with DAPI, which showed clear overlap with the F318L-Hog1-EGFP signal (Figure 4B).

**FIGURE 4.**
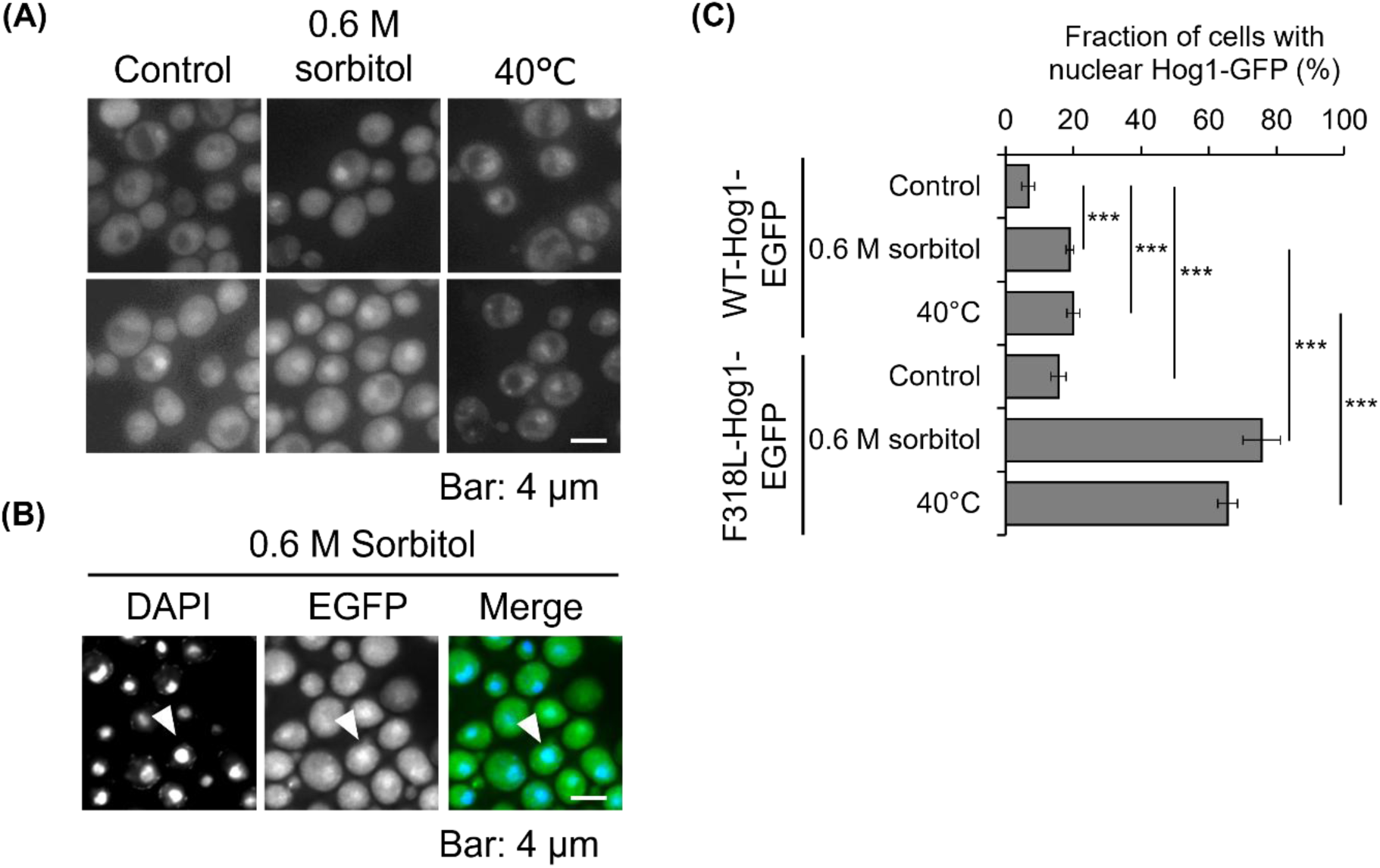
Hog1 subcellular localization under acute stress conditions. (A) Subcellular localization of WT-Hog1-EGFP and F318L-Hog1-EGFP under control conditions, osmotic stress (0.6 M sorbitol), and high-temperature stress (40 °C). Representative fluorescence images are shown. Scale bar, 4 μm. (B) Confirmation of nuclear localization of F318L-Hog1-EGFP under osmotic stress. Cells expressing F318L-Hog1-EGFP were stained with DAPI and observed by fluorescence microscopy. Arrowheads indicate nuclei with overlapping EGFP and DAPI signals. Scale bar, 4 μm. (C) Quantification of Hog1 nuclear localization under control conditions, osmotic stress (0.6 M sorbitol), and high-temperature stress (40 °C). The percentage of cells exhibiting nuclear-enriched Hog1-EGFP signal is shown. Data are the mean ± SD of at least three independent biological replicates. Statistical significance was determined by two-way ANOVA (main effect of condition, strain, and interaction); \*\*\**p* < 0.001

Similar to osmotic stress, high temperature (40 °C) promoted Hog1 nuclear localization in both strains; however, the extent of nuclear accumulation was again significantly greater for the F318L variant than for WT Hog1 (Figure 4A). Quantitative analysis of nuclear localization rates revealed that, whereas WT-Hog1-EGFP exhibited moderate stress-induced nuclear translocation, the F318L-Hog1-EGFP mutant showed significantly elevated nuclear localization under both osmotic and high-temperature stress conditions (Figure 4C). Collectively, these results indicate that the F318L mutation promotes enhanced and sustained nuclear accumulation of Hog1 in response to stress.

## 4 Discussion

This study has identified extracellular osmolarity as a determinant of yeast thermotolerance. High-temperature stress has been traditionally interpreted primarily in terms of intracellular molecular damage, particularly protein denaturation and aggregation (Hänninen et al. 1999; Trotter et al. 2001; Yost et al. 1991; Shahsavarani et al. 2012; T. D. Ingolia & Craig 1982). However, our findings now indicate that growth at high temperature is also mediated by the physicochemical properties of the extracellular environment. We have demonstrated that extracellular osmolarity increases during growth (Figure 2A, B), that the composition of solutes released changes at high temperature (Figure 2C-E), and that high extracellular osmolarity attenuates growth inhibition due to high temperature across phylogenetically diverse yeast species (Figure 1B-F).

From a physical perspective, these observations extend earlier reports of thermotolerance through osmotic supplementation (Costa et al. 2014; Papouskova & Sychrova 2007). Although ethanol production probably contributes to the increase in extracellular osmolarity during fermentation (Figure S2) (Zampar et al. 2013; Guijarro & Lagunas, 1984; Brauer et al. 2005; D’Amore et al. 1988), the reduction in glucose consumption that we observed at high temperature would be expected to decrease ethanol production. Nevertheless, extracellular osmolarity still increased under these conditions, suggesting that additional solutes are released during high-temperature growth. Elevated temperatures are known to affect plasma membrane permeability in yeast cells (Godinho et al. 2020; Guyot et al. 2015). In previous studies, moderately high temperatures such as 40 °C generally induced only limited changes in permeability; however, the prolonged cultivation used in this study (15 h) may have promoted gradual membrane destabilization. The precise composition of the extracellular osmolytes remains to be determined and represents an important subject for future investigation.

High temperature imposes not only proteotoxic stress, but also mechanical and osmotic disturbances. Yeast cells normally maintain a turgor pressure of ∼0.5–1.5 MPa by sustaining intracellular osmolarity above extracellular levels (Schaber et al. 2010; Mishra et al. 2022). Excessive intracellular osmotic pressure, however, increases water influx and may place mechanical stress on the plasma membrane and cell wall. Based on these established principles, our observations may be interpreted within a physical framework where changes in extracellular osmotic pressure influence cellular responses to high-temperature conditions. In this context, high extracellular osmolarity is expected to reduce the osmotic gradient across the cell boundary, potentially helping to mitigate temperature-induced stress. Although turgor pressure itself was not directly measured in this study, its involvement is inferred from the consistency of our observations with previous research.

In agreement with this hypothesis, deletion of the glycerol channel Fps1 resulted in pronounced heat sensitivity that was rescued by osmotic supplementation (Figure 1F). Moreover, as in previous studies (Dunayevich et al. 2018), the extracellular glycerol concentration increased under high-temperature conditions (Figure 3B), suggesting enhanced solute efflux. Recent studies have demonstrated that high pressure causes an increase in turgor pressure via rapid water influx, followed by activation of Fps1-mediated efflux of glycerol to prevent cell rupture (Mochizuki et al. 2023). These findings support a model in which high-temperature stress elevates turgor pressure, requiring solute efflux. This model is also consistent with a study on temperature changes and intracellular water potential (Watson et al 2023) showing that temperature elevation increases the ratio of free water to structured water, resulting in thermodynamic behavior similar to hypotonic treatment. Importantly, this physical buffering mechanism complements rather than replaces molecular damage-centric models of thermotolerance. Protein misfolding and aggregation remain central components of high-temperature stress, but our results indicate that physical stress derived from osmotic imbalance constitutes an additional dimension of cellular damage.

In parallel with extracellular osmotic modulation, intracellular accumulation of glycerol emerges as a critical intracellular protective mechanism. Glycerol stabilizes proteins and membranes and influences intracellular hydration states (Meng et al. 2004; Panadero et al 2006; Gekko et al. 1981; Pocivavsek et al. 2011)—properties that are particularly relevant under conditions promoting protein unfolding. We found that high-temperature and hyperosmotic conditions independently increased intracellular glycerol levels, and their effects were approximately additive (Figure 3A). Combining the two conditions resulted in intracellular glycerol concentrations that were substantially higher than those observed under either condition alone. Such pronounced accumulation suggests that glycerol may contribute not only to osmotic balance and protein stabilization, but also to the physicochemical state of the cytoplasm. As mentioned in the Introduction, yeast cells counteract temperature-dependent decreases in cytoplasmic viscosity (Longsworth 1954) through the accumulation of storage carbohydrates such as trehalose and glycogen (Persson et al. 2020). Although we did not examine the contribution of glycerol to cytoplasmic viscosity directly, the high intracellular concentrations observed under combined thermal and osmotic stress raise the possibility that glycerol may also influence intracellular physicochemical properties under these conditions.

The approximately additive increase in intracellular glycerol levels observed under combined temperature and osmotic stress indicates that temperature-dependent and osmolarity-dependent glycerol accumulation are not solely mediated by a single shared pathway. Consistent with previous observations (Siderius et al. 2000), glycerol accumulation during prolonged high-temperature cultivation occurred even in the absence of *HOG1* (Figure S4). Thus, sustained glycerol accumulation under thermal stress is regulated, at least in part, independently of canonical osmotic stress signaling. In contrast, the strain expressing the Hog1^F318L^ variant showed enhanced basal glycerol levels and improved growth at high temperature (Figure 3C, D). Shiraishi et al. (2018) observed the sequestration of Hog1 into stress granules at high temperature in *Candida boidini* and *S. pombe*, supporting a model in which HOG signaling does not contribute to glycerol accumulation during prolonged high-temperature cultivation. Given the above, the HOG signaling pathway seems to function as a regulatory component to amplify glycerol synthesis in acute stress but is not a primary determinant of long-term high-temperature adaptation. However, the regulatory factors that increase glycerol accumulation under prolonged high-temperature conditions remain unknown.

Taken together, our findings support a multilayered model of thermotolerance in which extracellular and intracellular osmotic regulation operate in concert. High temperature seems to increase intracellular osmolarity and promote solute efflux, thereby preventing excessive intracellular osmotic pressure. High extracellular osmolarity, in turn, buffers excessive turgor pressure and reduces mechanical stress. Simultaneously, intracellular glycerol accumulation stabilizes macromolecular structures and contributes to maintaining proteostasis under high-temperature stress. Thermotolerance thus emerges not only from intracellular damage repair systems, but also from the coordinated regulation of physicochemical parameters across the cell boundary. This cell–environment coupling perspective extends current models centered on protein quality control and osmotic stress signaling.

From an applied perspective, these findings present new strategies for engineering thermotolerant strains. Rather than focusing solely on enhancing the protein homeostasis network, approaches that regulate extracellular osmotic pressure or promote controlled glycerol accumulation might increase robustness under high-temperature fermentation. Such approaches challenge the conventional understanding of fermentation inhibition by high osmotic pressure; therefore, validation under conditions closer to actual fermentation production environments will be required.

This study positions extracellular osmolarity and turgor regulation as central components of yeast thermotolerance. By integrating physical buffering with biochemical protection, yeast cells can achieve sustained growth at high temperature. Future work should directly quantify turgor pressure dynamics, characterize extracellular metabolite composition, and elucidate the regulatory networks underlying HOG signaling-independent glycerol accumulation. Clarifying how mechanical forces intersect with metabolic and signaling pathways will advance understanding of stress adaptation derived from cell-environment interactions.

## Acknowledgements

The authors thank the National BioResource Project (NBRP), NITE Biological Resource Center, National Research Institute of Brewing, Japan, American Type Culture Collection for providing yeast strains. This work was supported by grants for a Grant-in-Aid for Young Scientists (JSPS KAKENHI Grant Number JP 24K17819) from the Japan Society for the Promotion of Science (JSPS) to Y.Y., and a Grant-in-Aid for Young Scientists (Y-2024-2-015) from the Institute for Fermentation, Osaka, to Y.Y.

## Data Availability Statement

The data that support the findings of this study are available from the corresponding author upon reasonable request.

## Author Contributions

Satoshi O., K.S., F.A., and Y.Y. conceived the study and designed the experiments. Saki O., R.I., A.A., K.K., Satoshi O., K.S., T.T., and Y.Y. performed the experiments. Saki O, Satoshi O., K.S., F.A., and Y.Y. analyzed the data and wrote the manuscript. All authors read and approved the final manuscript.

## Supporting information

Additional supporting information can be found online in the Supporting Information section.

**Figure S1:** Phylogenetic tree of yeast species based on the D1/D2 domain of the large subunit rRNA gene.

**Figure S2:** Temporal changes in extracellular osmolarity during cultivation at 30 °C.

**Figure S3:** Temperature-dependent relationship between optical density and dry cell weight.

**Figure S4:** Intracellular glycerol levels in *hog1*Δ cells cultivated at different temperatures.

