## Supplemental FIGURE S1-S4 for "High extracellular osmolarity promotes yeast thermotolerance through osmotic modulation and glycerol-dependent adaptation"

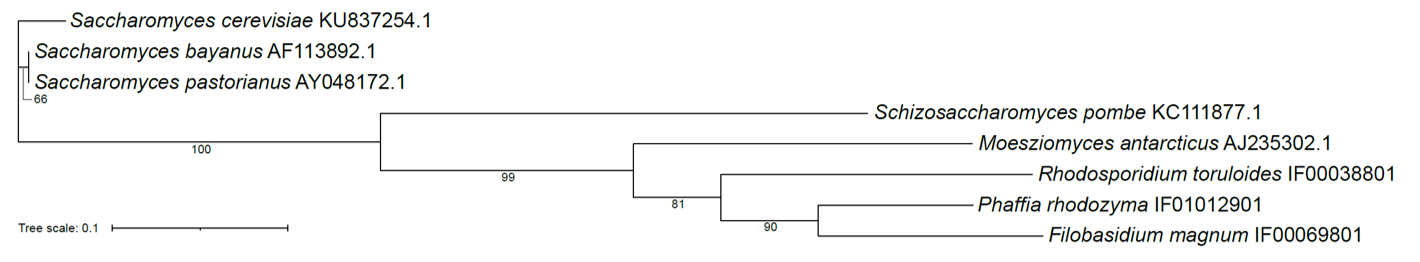

**FIGURE S1. Phylogenetic tree of yeast species based on the D1/D2 domain of the large subunit rRNA gene.**
Sequences were aligned using MAFFT, and poorly aligned terminal regions were trimmed to retain only positions shared by all taxa. Phylogenetic reconstruction was performed using the maximum likelihood method implemented in IQ-TREE 3.0.1 with ModelFinder for substitution model selection. Branch support was evaluated by ultrafast bootstrap analysis with 1000 replicates. Bootstrap support values greater than 60% are indicated at the corresponding nodes. Branch lengths represent the estimated number of nucleotide substitutions per site; scale bar corresponds to 0.1 substitutions per site.

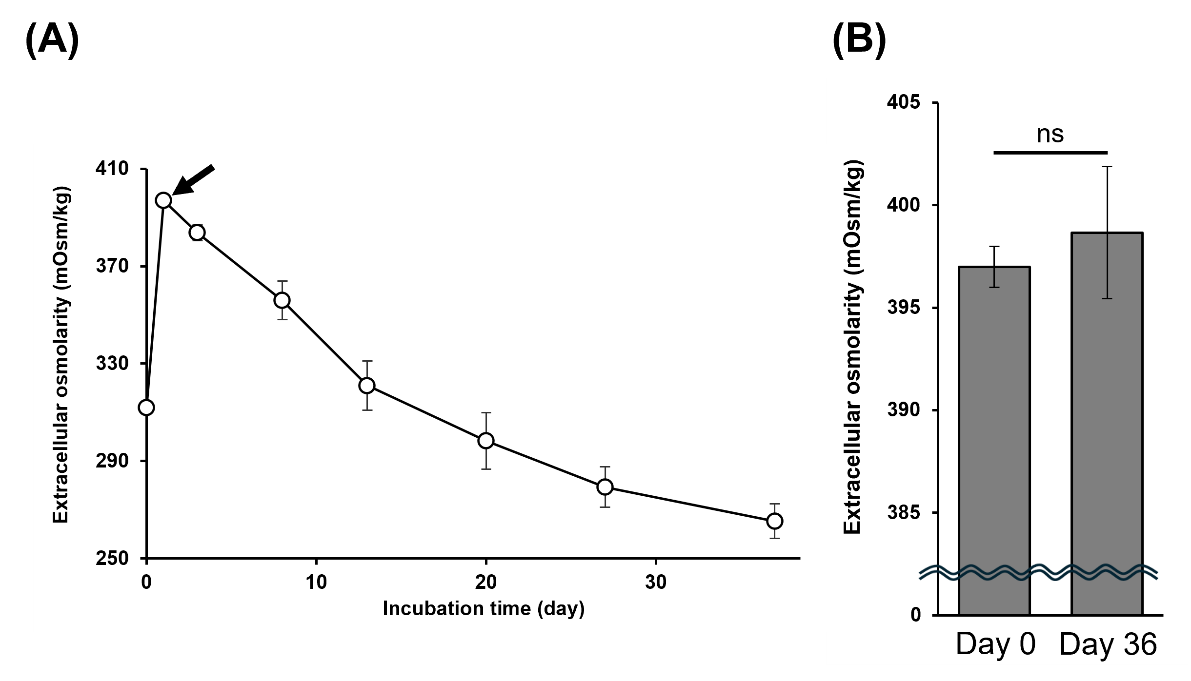

**FIGURE S2. Temporal changes in extracellular osmolarity during cultivation at 30 °C.**

(A) Extracellular osmolarity of the culture medium was monitored over time during growth at 30 °C. Arrows indicate the sampling times used for the corresponding cell-free incubation shown in (B). (B) Extracellular osmolarity measured at the indicated sampling times and after prolonged incubation (36 days) under identical conditions in the absence of cells. Data are the mean ± SD of at least three independent biological replicates. Statistical significance is indicated (ns, not significant).

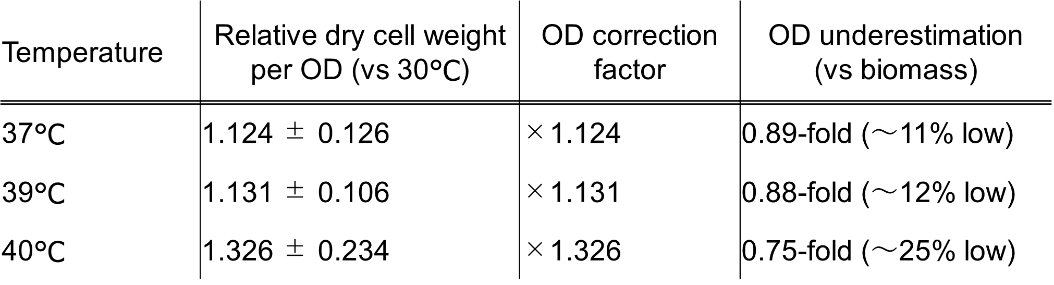

**FIGURE S3. Temperature-dependent relationship between optical density and dry cell weight.**

To assess the temperature dependence of biomass estimation based on optical density (OD_600_), the relationship between OD_600_ and dry cell weight (DCW) was examined at 30, 37, 39, and 40 °C. For each temperature, data are the mean ± SD of 10 or more independent biological replicates. Values are presented as relative DCW per OD_600_ normalized to the 30 °C condition. The resulting temperature-specific correction factors were used for OD_600nm_-based normalization throughout this study, including analyses of intracellular and extracellular glycerol levels (Figure 3).

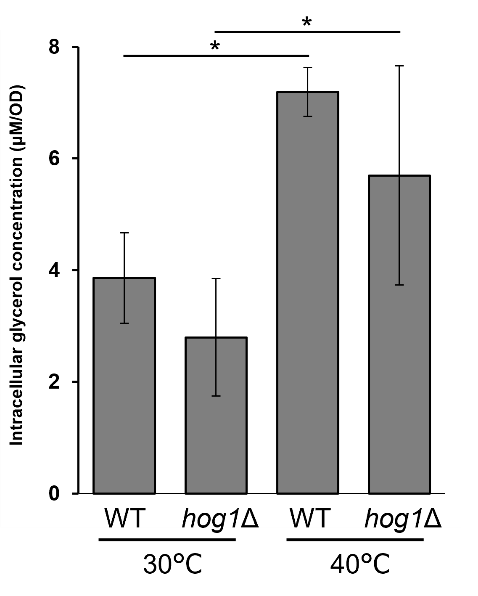

**FIGURE S4. Intracellular glycerol levels in *hog1*Δ cells cultivated at different temperatures.**
Wild-type and *hog1*Δ cells were cultivated at optimal or high temperature under the indicated conditions. Intracellular glycerol levels were quantified after cultivation and normalized to cell density. Data are the mean ± SD of at least three independent biological replicates. Statistical significance was determined by two-way ANOVA (main effect of temperature); **p* < 0.05.
